# White matter geometry confounds Diffusion Tensor Imaging Along Perivascular Space (DTI-ALPS) measures

**DOI:** 10.1101/2025.04.14.648733

**Authors:** Kurt G Schilling, Allen Newton, Chantal Tax, Markus Nilsson, Maxime Chamberland, Adam Anderson, Bennett Landman, Maxime Descoteaux

## Abstract

**Introduction:** The perivascular space (PVS) is integral to glymphatic function, facilitating fluid exchange and waste clearance in the brain. Diffusion Tensor Imaging Along the Perivascular Space (DTI-ALPS) has been proposed as a non-invasive marker of perivascular diffusion, yet its specificity remains unclear. ALPS measures assume that radial asymmetry in white matter diffusivity predominantly reflects PVS contributions. However, anatomical and microstructural confounds may influence these metrics.

**Methods:** We systematically evaluated potential biases in ALPS-derived measures using high-resolution, multi-shell diffusion MRI from the Human Connectome Project (HCP) and high-field imaging. Specifically, we examined (1) the prevalence of radial asymmetry across white matter, (2) the influence of crossing fibers on ALPS indices, (3) the impact of axonal undulations and dispersion, and (4) the spatial alignment of vasculature with white matter in ALPS-associated regions.

**Results:** Radial asymmetry is widespread across white matter and persists even at high b-values, suggesting a dominant contribution from axonal geometry rather than faster PVS-specific diffusion. Crossing fibers significantly inflate ALPS indices, with greater radial asymmetry observed in regions with a greater prevalence of crossing fibers. Furthermore, anisotropic axonal dispersion and undulations introduce systematic asymmetry independent of perivascular diffusion. Finally, high-resolution vascular imaging reveals substantial heterogeneity in medullary vein orientation, challenging the assumption that PVS consistently aligns with the left-right axis in ALPS regions.

**Conclusion:** ALPS indices are significantly influenced by white matter microstructure, including fiber crossings, undulations, and dispersion. These findings suggest that ALPS-derived metrics may not provide a direct measure of glymphatic function but rather reflect underlying axonal geometry. Interpretations of ALPS-derived metrics as biomarkers of glymphatic function must consider these anatomical complexities, and future studies should integrate advanced modeling approaches to disentangle perivascular contributions from white matter structure.

## Introduction

The perivascular space (PVS), also known as the Virchow-Robin space, is a key anatomical structure facilitating fluid exchange between cerebrospinal fluid (CSF) and interstitial fluid along the brain’s vascular pathways [1]. This fluid movement is integral to the glymphatic system, a recently characterized network implicated in the clearance of metabolic waste, including amyloid-beta and other neurotoxic solutes from the brain parenchyma [2]. Disruptions in glymphatic function have been linked to neurodegenerative diseases such as Alzheimer’s and Parkinson’s disease [3], highlighting the need for robust, non-invasive imaging methods to assess perivascular dynamics.

Diffusion Tensor Imaging Along the Perivascular Space (DTI-ALPS) has recently emerged as a promising, non-invasive approach for probing glymphatic function [4]. First introduced by Taoka et al. in 2017, the DTI-ALPS method utilizes diffusion tensor imaging (DTI) to measure water diffusivity along the PVS, leveraging specific regions-of-interest (ROIs) adjacent to the lateral ventricles where white matter tracts and medullary veins intersect at near-orthogonal angles. In principle, this unique geometry allows the isolation of diffusion signals attributed to perivascular flow, culminating in a single scalar measure – the ALPS-index – that aims to serve as an indirect marker for glymphatic function.

The appeal of the ALPS-index lies in its ease of application, requiring only standard DTI acquisitions and minimal computational effort. Because of this, it has been rapidly adopted across numerous studies with the aim to investigate glymphatic function in both health and disease [5]. For example, the ALPS-index has been correlated with disability and cognitive performance in Alzheimer’s disease [6, 7], Parkinson’s disease [8, 9], and demyelinating diseases [10, 11], while also demonstrating sensitivity to aging and other neurophysiological changes [7, 8, 12, 13]. Despite its growing application, the specificity of the ALPS-index for perivascular diffusion remains unclear, as various microstructural and anatomical factors may contribute to the measured signal.

A key assumption of the ALPS-index is that diffusion within and around white matter tracts exhibits radial symmetry, specifically that the second and third eigenvalues of the diffusion tensor (λ2 and λ3) are equal. Recent work by Wright et al. [14], has demonstrated that radial asymmetry (λ2 > λ3) is not confined to ALPS-specific regions but is widespread throughout the brain. This observation suggests that ALPS-index may not solely reflect perivascular diffusion but also reflects axonal contributions [14]. We first aim to reproduce and validate these findings, hypothesizing that radial asymmetry exists broadly throughout the white matter and persists at diffusion weightings that are more sensitive to axonal contributions (i.e., high b-values), implicating neurite-related effects. At high b-values, diffusion-weighted imaging strongly attenuates signal from compartments with rapid and less-restricted diffusion, such as the extra-axonal space and perivascular spaces (PVS), isolating the contribution from the more restricted intra-axonal compartment. PVS typically occupies spaces on the order of 10–45 μm in height adjacent to vessels [15], substantially larger than the narrow calibers of axons (∼1 μm), reinforcing the expectation that high b-value signal predominantly reflects axonal microstructure.

A potential confounding factor is the presence of crossing fibers, a well-documented challenge in diffusion MRI. White matter of the brain is often comprised of multiple fiber populations intersecting within a single voxel, an effect that occurs in anywhere from ∼70-90% of all white matter voxels [16, 17], complicating the interpretation of any diffusion-derived metrics. Georgiopoulos et al. [18] recently highlighted that crossing fibers significantly inflate the ALPS-index (i.e., increasing the ratio of λ2/λ3), suggesting a lack of specificity for perivascular diffusion and highlighting an axonal contribution to this index. Our second aim is to further investigate the effects of crossing fiber confounds on the ALPS-index, hypothesizing not only that this measure is influenced by crossing fibers, but that radial asymmetry exists even in voxels with single-fiber populations.

Another possible confounding factor is the presence of axonal undulations, a phenomenon characterized by wave-like deviations along the course of axonal fibers. These undulations have been observed in histological preparations throughout various white matter regions [19–22], and are believed to serve a protective role, allowing axons to accommodate mechanical stress without sustaining damage [23]. Undulations have been extensively studied in the diffusion MRI community [24–28]. Several studies have compared the diffusion properties of straight versus undulating axons, finding that undulation can induce directional-dependent diffusivity even in single-fiber regions [27].

This phenomenon, where undulations in one plane can increase diffusivity while leaving the orthogonal plane unaffected, can introduce radial asymmetry independent of perivascular diffusion, further confounding the interpretation of the ALPS-index. Thus, our third aim was to test the hypothesis that radial asymmetry persists in single-fiber regions, and can be attributed to axonal undulation effects.

Finally, the ALPS-index assumes that the vasculature (and hence the PVS) runs consistently in the left-right (x) direction of the brain and is orthogonal to the white matter tracts of the superior-inferior (z) projection pathways and the anterior-posterior (y) association pathways [4]. Although medullary veins running in the left-right direction lateral to the ventricles has been consistently observed in several studies, variability in vascular orientation has also been reported [29–33], challenging the robustness of these assumptions. Our fourth aim is to assess whether individual variability in vascular structure and geometry results in inconsistent alignment of vasculature along the left-right axis, and whether white matter tracts in ALPS-related regions are reliably orthogonal to vasculature, thereby testing key assumptions underlying the ALPS methodology.

To systematically evaluate these confounds, we leveraged a high-resolution, high-quality diffusion imaging dataset to test each hypothesis. We first investigate the prevalence of radial asymmetry throughout the brain and ask whether this asymmetry occurs with different diffusion weightings. Second, we investigate the effects of crossing fibers on radial asymmetry (and consequently, the ALPS-index), and ask whether this phenomenon exists even within regions that do not contain crossing fibers. Third, we investigate the effects of axonal undulation and dispersion on radial asymmetry measures and use multi-compartment modeling to ask whether this explains asymmetry, again even in single fiber regions. Fourth, we use high-resolution, high-field imaging to make direct comparisons of vasculature and white matter orientation in the same subjects, asking whether ALPS-associated vasculature is indeed oriented right-to-left and perpendicular to white matter pathways, and where this assumption is (or is not) violated. Finally, given previous reports of APLS-index sensitivity to aging, we assessed whether these confounding factors also exhibit age-related changes, which would further support an axonal contribution to the ALPS signal. Overall, this study aims to refine the interpretation of the ALPS-index as a biomarker of glymphatic function.

## Methods

### Prevalence of Radial Asymmetry

First, we examine the prevalence of radial asymmetry across the brain and its dependence on diffusion weighting. We do this using two open-source datasets, with the goal to measure radial asymmetry in ALPS-specific ROIs as well as other ROIs throughout the cerebral white matter, and at multiple diffusion weightings.

#### Datasets

The datasets used in this study come from the Human Connectome Project [34], which aims to map the structural connections and circuits of the brain and their relationships to behavior by acquiring high-quality magnetic resonance images. We used diffusion MRI data from the Human Connectome Project Young Adult (HCP-YA) and the Human Connectome Project Aging (HCP-A) study. The Young Adult cohort was composed of a subset of N=105 participants aged 21 to 35 years. The Aging cohort was composed of subset of N=50 participants aged 35 to 90 years. The diffusion MRI acquisitions were slightly different for each dataset and tailored towards the population under investigation. For the Aging cohort, a multi-shell diffusion scheme was used, with b-values of 1500 and 3000 s/mm2, sampled with 93 and 92 directions, respectively (24 b = 0). The in-plane resolution was 1.5 mm, with a slice thickness of 1.5 mm. For the Young Adult cohort, the minimally preprocessed data [35] from Human Connectome Projects (Q1-Q4 release, 2015) were acquired at Washington University in Saint Louis and the University of Minnesota [34] using a multi-shell diffusion scheme, with b-values of 1000, 2000, and 3000 s/mm2 ,sampled with 90 directions each (18 b = 0). The in-plane resolution was 1.25 mm, with a slice thickness of 1.25 mm. For all diffusion data, susceptibility, motion, and eddy current corrections were performed using TOPUP and EDDY algorithms from the FSL (v6.0.7.9) package following the minimally preprocessed HCP pipeline [35]. Structural images with T1-weighting for all cohorts were acquired with an MPRAGE sequence, with a resolution of 0.8 mm isotropic for the Aging cohort, and resolution of 0.7 mm isotropic for the Young Adult cohort.

#### Radial Asymmetry

For each subject, the diffusion tensor [36, 37] was calculated voxel-wise (using an iterated weighted least-squares algorithm from MRtrix3 [38]), from which the primary (λ1), secondary (λ2), and tertiary (λ3) eigenvalues were derived. The tensor was derived for each diffusion weighting (b-value) separately. Following [14], radial asymmetry was calculated as the ratio of the secondary to tertiary eigenvalues (λ2/ λ3) as a measure of asymmetry orthogonal to the dominant direction of diffusion.

#### Regions of Interest

The Johns Hopkins University (JHU) white matter atlas [39] was used to define 35 white matter ROIs. These regions represented locations of major association, projection, and commissural pathways of the brain. In addition to these, we used 4 ALPS-specific regions designed for automated computation of the ALPS-index [40] manually delineated on the JHU-ICBM-FA template (and available at https://github.com/gbarisano/alps). These regions represent the projection and association fibers at the level of the lateral ventricle body within the superior corona radiata (left and right SCR) and within the superior longitudinal fasciculus (left and right SLF) and were defined as spheres with 5mm diameters. We note that these are sub-sets of the much larger JHU ROIs bearing the same name but intended to isolate the regions where vasculature runs left-to-right and is orthogonal to the projection and association fibers. Within each region we derive the average asymmetry. Additionally, we calculate a measure of coherence of the second eigenvector (V2 Coherence) to assess whether axial asymmetry is due to noise (which would result in a low coherence) or vascular/axonal geometry (which would result in a non-negligible coherence). This was calculated as the fractional anisotropy of the orientation tensor derived by averaging the outer products of all secondary eigenvectors within each region.

### Crossing Fibers

Second, we investigated the influence of crossing fibers on ALPS-related measures. We performed a simulation to investigate crossing fiber confounds on radial asymmetry, and empirically measured the relationship between features of crossing fibers and radial asymmetry.

#### Simulations

We simulated crossing fibers by generating two fiber populations crossing at a specific crossing angle, and with a specific crossing fraction, following [41]. The crossing angle was defined as the angle between the two fiber populations (ranging from 0-90°), while the crossing fraction was defined as the fraction of the secondary fiber population (ranging from 0-0.5). Simulations were performed using a multi-tensor signal simulation (1 ≤ L1 ≤ 2 mm2/s; 0.1 ≤ L2=L3 ≤ 0.6 mm2/s) for crossing angles from 0-90° in steps of 1°, and crossing fraction from 0-0.5 in steps of 0.01, with 1000 crossing fiber voxels simulated for every geometric configuration. Noise was added in quadrature (i.e., Rician noise) at a level matching that derived from HCP datasets (SNR∼20-30), the diffusion tensor was calculated, from which radial asymmetry derived.

#### Single and Crossing Fibers

We derived equivalent voxel-wise measures of crossing angle and crossing fraction from HCP and HCPA data using MRtrix3 functions. This procedure included (1) estimating response functions using dwi2response dhollander algorithm [42, 43] (2) performing multi-shell multi-tissue spherical deconvolution to estimate the fiber orientation distribution (FOD) using the dwi2fod msmt_csd [42, 43] function (3) deriving peaks of the FOD using sh2peaks function, and (4) calculating the number of peaks that have a crossing fraction >0.05% of the maximum peak value using mrcalc + mrmath functions. From this, for each region of interest, we calculated the single fiber fraction (the fraction of voxels containing only a single fiber population), the crossing angle (the angle between the two maximum peaks, as in the simulated data), and the crossing fraction (the total fraction of crossing fibers, as in the simulated data). Finally, derived the asymmetry measure in each ROI (both JHU ROIs and the 4 ALPS-Specific ROIs) as described in the *Prevalence of Radial Asymmetry* Section.

### Undulation

Third, we assessed the role of axonal undulations in introducing asymmetry. This again included simulations and empirical measurements of radial asymmetry obtained from biophysical modeling.

#### Simulations

Following Nilsson et al. [27], we simulated axonal undulations using a Monte Carlo framework in which axons were modeled as sinusoidal curves defined by three parameters: the axon diameter (d), the undulation wavelength (L), and the undulation amplitude (A). Briefly, we ran simulations in which water molecules executed random walks within the confines of these undulating geometries, with reflective boundaries representing the axonal membranes. Specifically, we considered two scenarios: one mimicking microscale undulations with d = 1, L = 24, and A = 4, and another representing macroscale undulations with d = 1, L = 300, and A = 50. Simulations were run using the diffusion encoding protocol of the HCP data used in the current study. These settings allowed us to generate diffusion propagators and signal attenuation curves comparable to those measured in diffusion MR experiments. From these, we fit a diffusion tensor and calculated the asymmetry index (λ2/ λ3) both before and after noise was added to the signal.

#### Modeling anisotropic orientation dispersion

Building on our simulation findings, we analyzed empirically acquired diffusion MRI data by applying a multi-compartment (i.e., biophysical) model of tissue microstructure that explicitly represents the orientation distribution and dispersion of neurites. Following Tariq et al. [44] , we used our data to fit the Neurite Orientation Dispersion and Density Imaging (NODDI) model, which describes neurite orientations using one of two distributions: the Watson distribution, which assumes isotropic dispersion around a dominant orientation, and the Bingham distribution, which accounts for anisotropic dispersion by separately capturing the spread along the primary and secondary dispersion axes.

To determine which distribution best explained the observed diffusion MRI signal, we performed a model comparison using the Bayesian Information Criterion (BIC). This approach allowed us to assess whether the anisotropic (Bingham, i.e., radially asymmetric) or isotropic (Watson, i.e., radially symmetric) formulation provided a more parsimonious explanation of the data, with lower BIC values indicating better model fit. Finally, for the Bingham-distribution model, we estimated the dispersion anisotropy index (DAB) - a metric quantifying the degree of anisotropy in neurite orientation dispersion.

It is important to note that while both the Bingham and Watson models were fit to all voxels, our primary analysis focused on single-fiber voxels. We hypothesized that radial asymmetry (i.e., dispersion anisotropy) exists even in the absence of crossing fiber populations.

### Asymmetry and Age

Fourth, given prior reports linking ALPS-index to aging, we first confirmed radial asymmetry changes with age, then examined whether confounding factors exhibit age-related effects. If observed, this would further suggest axonal contributions. For this, we simply fit a linear regression model using age as the predictor variable and feature (radial asymmetry, V2 Coherence, Crossing Angle, Single Fiber Fraction, Crossing Fraction, Dispersion Anisotropy) as the response variable. This was performed globally over the entire white matter, as well as for each ROI independently (both JHU ROIs and the 4 ALPS-Specific ROIs), where the linear correlation coefficient and effect sizes were derived, and only shown if statistically significant at p<0.05 after false discovery rate correction.

### Geometrical configuration of vasculature and white matter

Finally, we perform a qualitative and quantitative evaluation of the orientation of vasculature throughout the white matter, and its relationship to white matter fiber orientation. This hypothesis was tested using a separate, high-resolution dataset (acquired at Vanderbilt University Medical Center, and approved by the Vanderbilt Institutional Review Board - IRB #020623) that allowed visualization, segmentation, and orientation derivation of both vasculature and white matter structures, allowing for a voxel-wise comparison in the same subjects (N=4).

#### Data Acquisition

High-resolution imaging data were collected using susceptibility-weighted imaging (SWI) for vascular visualization and diffusion-weighted imaging (DWI) for white matter fiber orientation estimation. SWI was performed on a 7T MRI scanner using a multi-echo 2D gradient-recalled echo sequence (0.2×0.2 resolution, 1mm slice thickness, N=5 averages). Phase images underwent high-pass filtering and a power transformation before being combined with magnitude images to enhance deoxyhemoglobin contrast. The resulting images were then super-resolved to a 0.2 mm isotropic resolution using the SMORE deep learning algorithm, followed by co-registration and averaging to optimize signal-to-noise ratio. DWI was acquired on a 3T scanner using a multi-shell acquisition with four b-values (b = 500, 1,000, 2,000, and 3,000 s/mm2). Data preprocessing included denoising (using both MPPCA [45] and Patch2Self [46]), and correction for susceptibility distortions, subject motion, and eddy current correction [47] using the PreQual preprocessing pipeline [48].

#### Vasculature Orientation Estimation and White Matter Orientation Estimation

Vascular structures were extracted from high-resolution SWI using an adaptive non-local means filter, followed by a Frangi vesselness filter to identify tubular structures. This segmentation provided voxel-wise vascular orientation estimates. To minimize partial volume contamination, white matter masks were applied, and segmented vasculature was downsampled to a 2 mm isotropic resolution for direct comparison with diffusion data. For white matter analysis, fiber orientation distributions (FODs) were estimated using multi-shell, multi-tissue spherical deconvolution. Up to three fiber orientations per voxel were extracted, corresponding to dominant white matter pathways.

#### Orientation Comparison

We made three primary orientation comparisons. First, we compared the angular difference between vasculature and V1, investigating the assumption that vasculature is orthogonal to WM in ALPS-ROIs. Second, we compared the angular difference between vasculature and V2, as V2 should align with vasculature if it was the primary determinant of diffusivity orthogonal to WM pathways. Third, we compared the angular difference between vasculature and the right-to-left (RL) orientation, investigating the assumption that the vasculature is oriented RL in ALPS-ROIs.

## Results

### There is widespread radial asymmetry in diffusivity across white matter

**Figure 1** shows the radial asymmetry index (defined as the ratio of the secondary to tertiary eigenvalues, λ2/λ3) across a representative HCP subject, for multiple sagittal, axial, and coronal slices (example HCP-A subject is given as **Supplementary Figure 1**).

**Figure 1.**
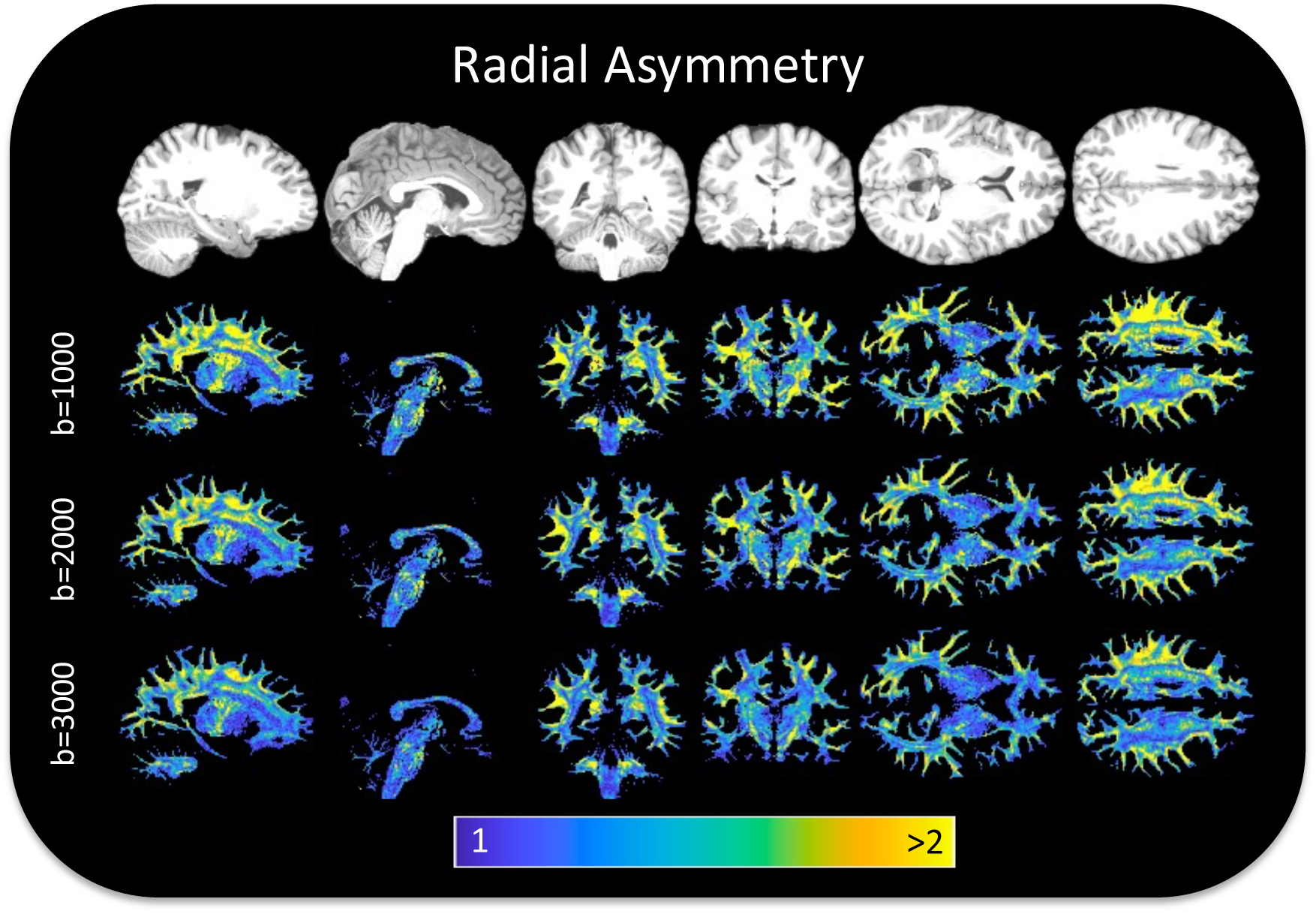
Radial asymmetry is widespread throughout white matter. Sagittal, coronal, and axial slices of an example HCP subject show radial asymmetry at all diffusion weightings, and throughout white matter, with most regions exhibiting average asymmetry values ∼1.3-1.8, with many voxels >2.

Radial asymmetry is prevalent throughout the entire brain, often greater than 1.5-2, meaning that the secondary eigenvalue is 1.5-2x greater than the tertiary eigenvalue. This asymmetry is observed throughout all b-values.

**Figure 2** confirms quantitatively that all white matter regions exhibit radial asymmetry in the HCP dataset (Results for HCP-A are shown in **Supplementary Figure 2**). Most regions exhibit average asymmetry values ∼1.3-1.8 (**Figure 2**, **top**), including the ALPS-specific indices, all of which are greater values than that expected by signal noise alone. The secondary eigenvectors are largely coherent throughout regions (**Figure 2**, **middle**), again, suggesting some coherent tissue (axonal, vascular) structure rather than noise. Finally, these values are observed at all b-values (**Figure 3**, **bottom**), even high b-values where PVS signals are expected to be absent (see Discussion: *Perivascular space size and shape and diffusion sensitivity*).

**Figure 2.**
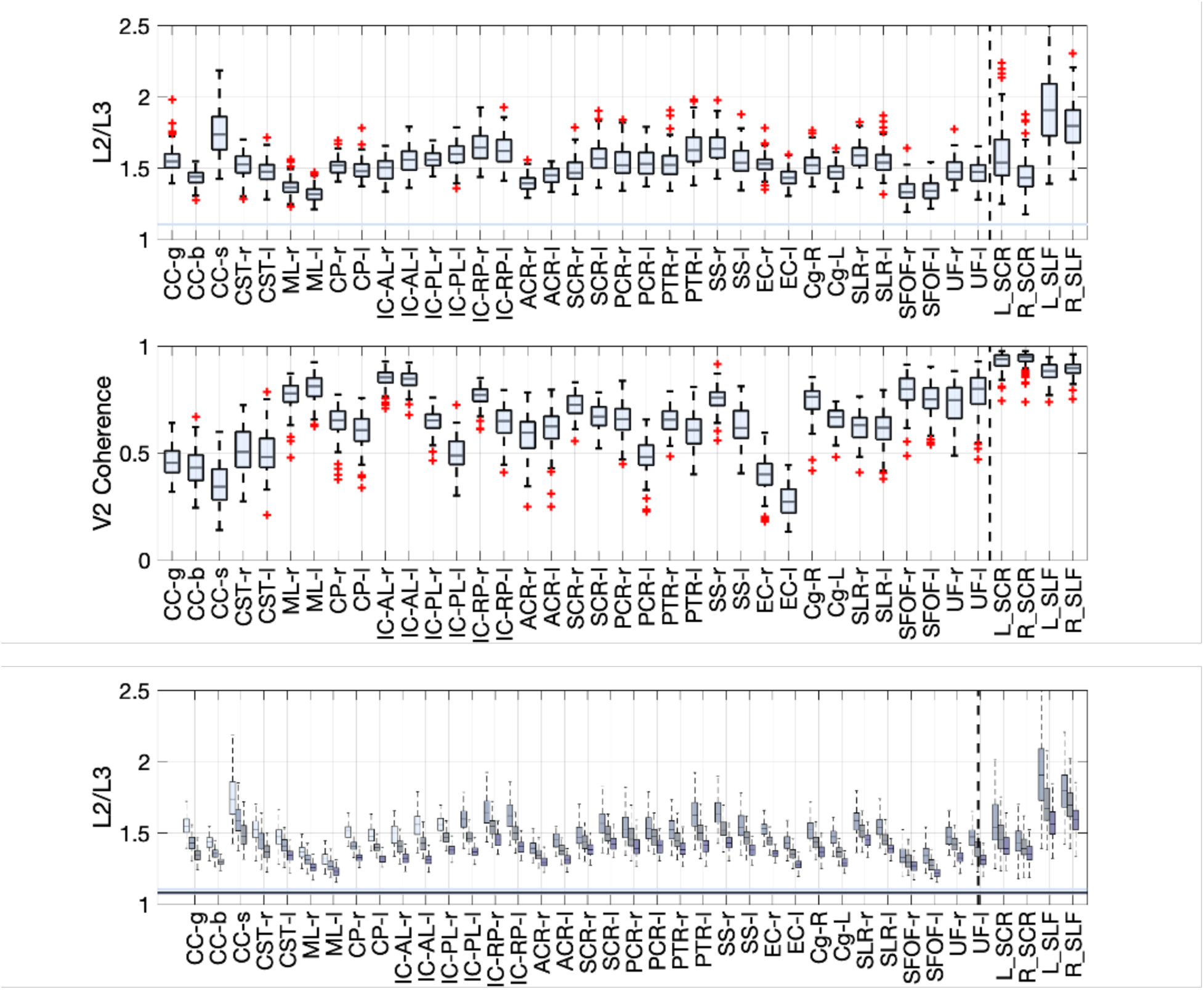
Most white matter regions exhibit radial asymmetry (“L2/L3”). Radial asymmetry (L2/L3) is greater than 1, and greater than noise (color bar) for all JHU white matter and ALPS-specific regions (top, b=1000 data)(note that the four ALPS-specific regions are the rightmost regions in each plot to the right of the dashed vertical line). The secondary eigenvector is coherent through these white matter regions (middle), suggesting this is not a noise-related effect. Radial asymmetry remains at all b-values, where b-values of 1000, 2000, 3000 s/mm2 are shown from light to dark for each region (bottom).

**Figure 3.**
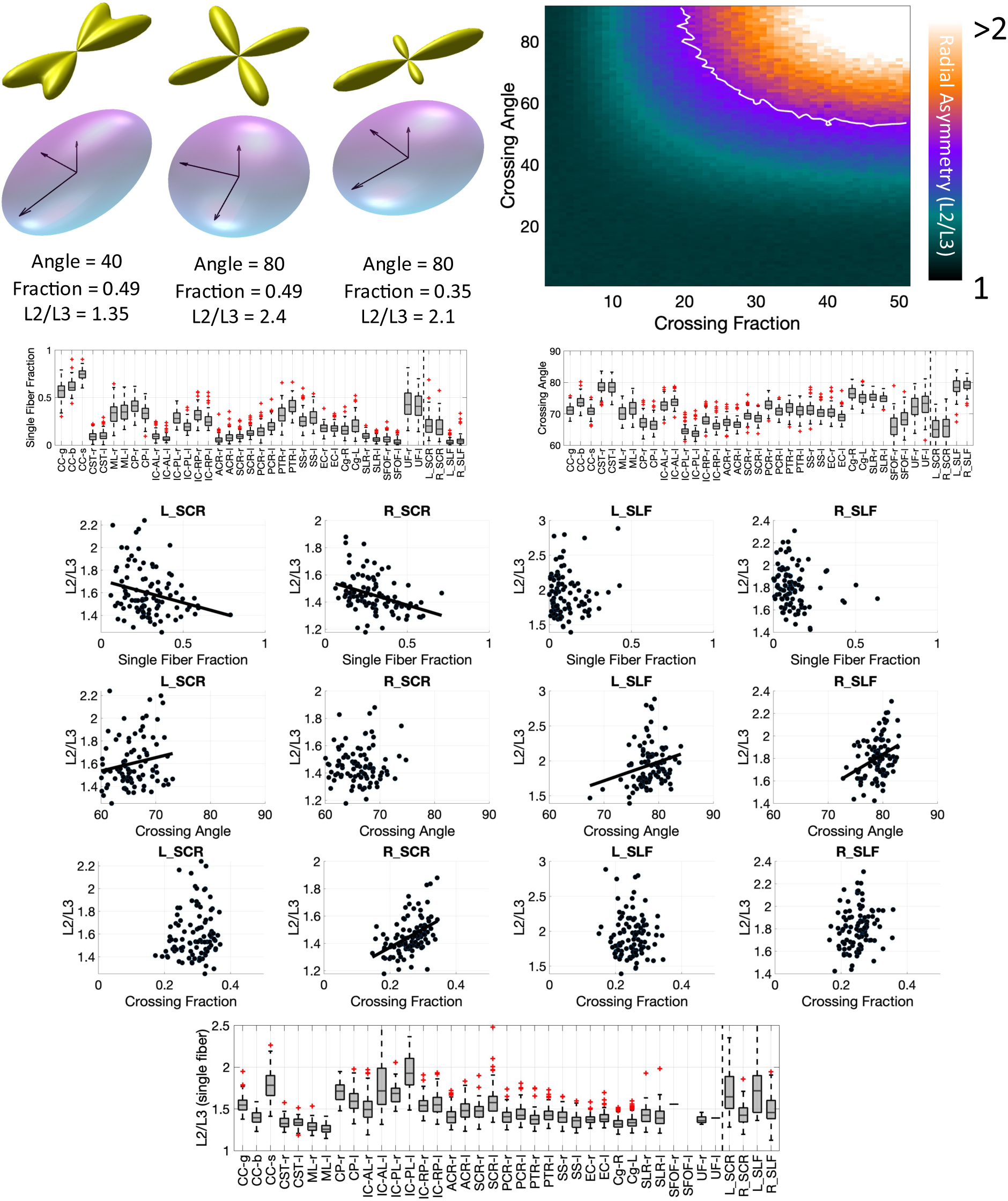
Most white matter regions contain crossing fibers, which contribute to radial asymmetry. (First Row) crossing fibers at different crossing angles and fiber fractions, resulting in radial asymmetry are visualized, as well as simulation results showing that increasing crossing angle and crossing fraction result in increased radial asymmetry. (Second Row) Single Fiber Fraction and Crossing Angle are shown averaged in each white matter region. (Middle) Radial asymmetry is plotted against Single Fiber Fraction, Crossing Angle, and Crossing Fraction for ALPS-specific ROIs, across subjects, confirming that crossing fibers significantly influence asymmetry measures. (Bottom) Axial asymmetry remains even in single fiber voxels within each region.

### Most white matter regions contain crossing fibers, which contribute to radial asymmetry

**Figure 3** shows exemplar crossing fiber glyphs (**Figure 3**, **top**), for two crossing fibers at varying angles and varying fiber fractions, as well as the derived diffusion tensor ellipsoid, eigenvalues, and asymmetry index. Intuitively, a greater crossing angle, and greater crossing fraction, lead to higher radial asymmetries. Our simulation experiments (**Figure 3**, **top**) confirm this, showing that orthogonal crossings and greater crossing fractions lead to radial asymmetry values frequently above 2. Here, the white line indicates the average asymmetry value observed in **Figure 2**, and suggests that most white matter contains crossing fibers, with not insignificant crossing fractions.

This is confirmed by extracting single fiber fraction and crossing angle across white matter regions (**Figure 3**, **second row**). Here, most regions (except for corpus callosum segments) have a very low single fiber fraction, with crossing angles generally between 60-90 degrees. Specifically within ALPS-related regions (**Figure 3**, **third row**), we observed that radial asymmetry increases as the single fiber fraction decreases, indicating greater asymmetry with more crossing fibers. Similarly, radial asymmetry also increases with larger average crossing angles, suggesting that both the number of crossing fibers and the angles at which they cross significantly contribute to asymmetry. However, even limiting analysis to single fiber regions, we find that radial asymmetry still exists throughout white matter (**Figure 3**, **bottom**), which suggests that crossing fibers alone do not explain asymmetry. Similar results are observed in HCP-A dataset, provided in **Supplementary Material**.

To further confirm that crossing fibers confound ALPS-related analysis, we show the ALPS-related ROIs in two example subjects in **Figure 4**, with FODs overlaid and voxels colored by the number of fiber populations. Most voxels in these regions contain crossing fibers (2 fibers or 3 fiber populations), with only a small minority unaffected by this effect. Visually, there is no region of this size that includes only single fiber voxels.

**Figure 4.**
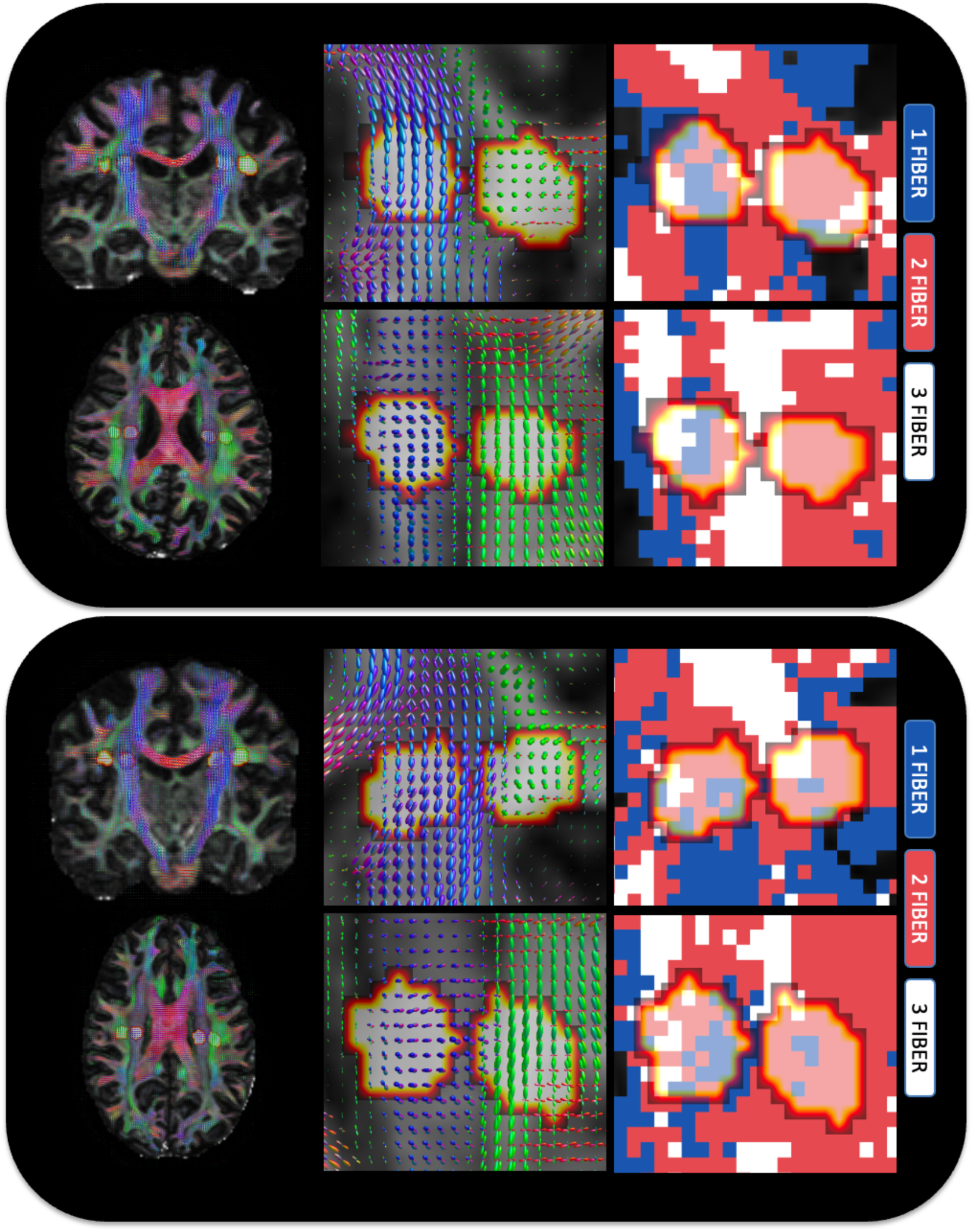
Crossing fibers are widespread throughout the white matter, and throughout ALPS-specific regions. For two example subjects, FODs are shown in coronal and axial slices, showing that many of these regions contain 2 or 3 fiber populations (i.e., are influenced by crossing fibers).

### Most white matter regions contain anisotropic dispersion and/or undulation

**Figure 5** shows example geometrical configurations that may cause axial asymmetry (**Figure 5**, **top**). This includes an undulation of amplitude A and length L within plane, or a dispersion of axons within the plane, both leading to similar diffusivity patterns with V1 (up and down) greater than V2 (left-right) and both greater than V3 (through-plane). Our Monte Carlo simulations of a microscopic undulations (A=4um, L=24um) and macroscopic undulations (A=50um, L=300um) confirm that both geometries can lead to asymmetries between 1.36-1.48 or 2.06-2.43, respectively, depending on b-value. These values are in-line with asymmetry observed in our data.

**Figure 5.**
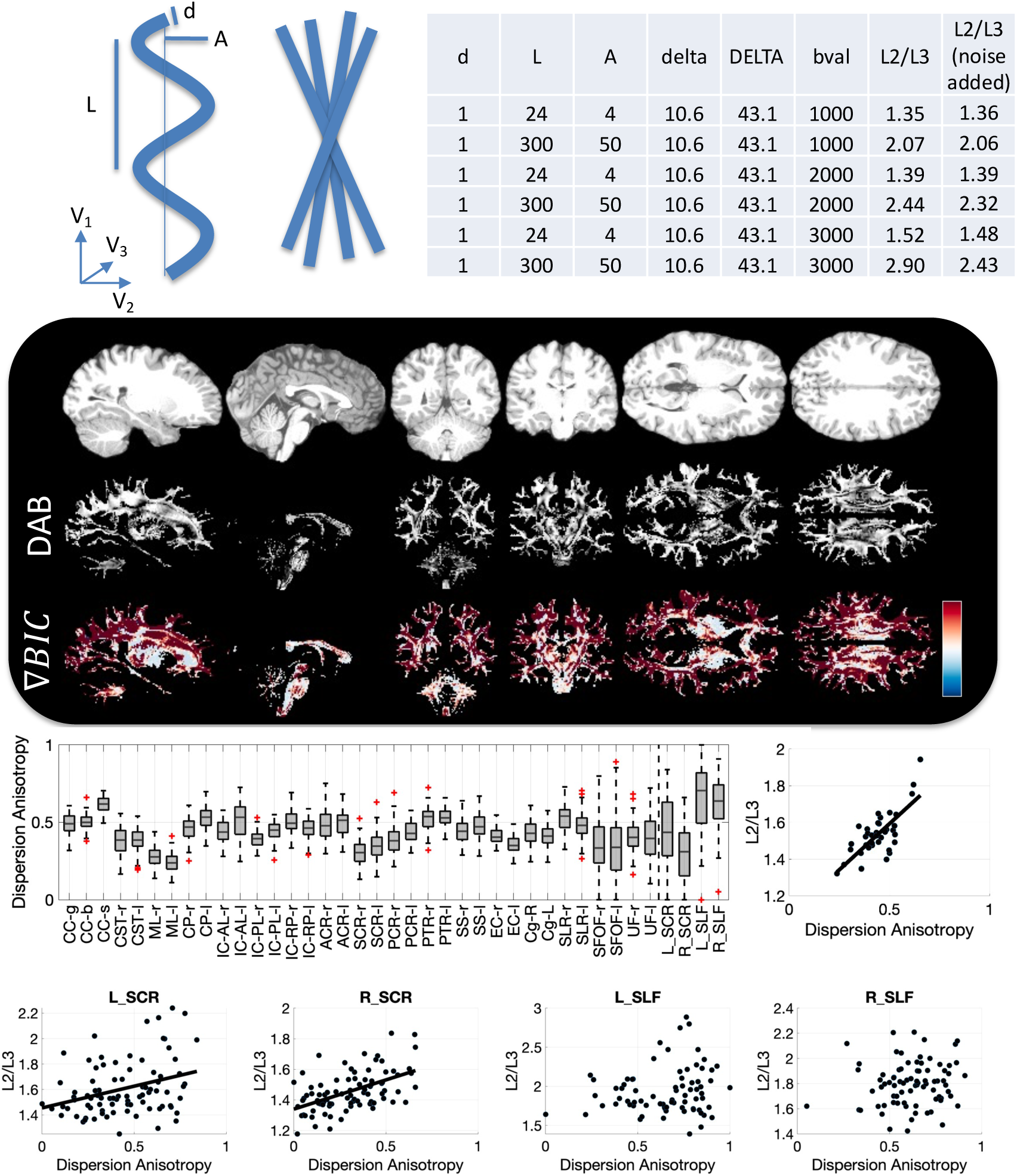
Dispersion and Undulation contribute to radial asymmetry. (Top) example undulation and dispersion cartoons, highlighting within-plane orientation dispersion. Simulation results with two substrates show that undulations can contribute to radial asymmetry on par with that observed in empirical data. (Second Row) Example sagittal, coronal, and axial slices showing the Dispersion Anisotropy Index (DAB) (gray-scale) and the difference in BIC between Watson and Bingham distributions (Red indicates preference for Bingham, Blue indicates preference for Watson distribution). (Third Row) Dispersion Anisotropy (DAB) across all white matter regions; radial asymmetry plotted against Dispersion Anisotropy (DAB) across all regions shows strong relationship between these two measures of asymmetry. (Bottom Row). Similarly, Radial Asymmetry plotted against DAB across subjects shows strong relationships between these two measures in ALPS-specific ROIs, with a trend-line shown for regions with statistically significant associations.

Results from NODDI model fitting are shown for an example subject (**Figure 5**, **second row**), where the dispersion anisotropy index (DAB) and BIC is shown for sagittal, coronal, and axial slices. DAB is high throughout the entire white matter, frequently >0.5, empirically. A positive BIC suggests a better model fit for a Bingham (radially asymmetry) over a Watson (radially symmetric) distribution. Here, positive or near-equal BIC is observed throughout the white matter. Together, these confirm that planar-dispersion or undulations exist (as well as crossing fibers).

Quantifying DAB in single fiber regions only (unaffected by crossing fibers) (**Figure 5**, **third row**), we see most regions have DAB ∼0.3-0.5, including the ALPS-specific regions, which match our qualitative observations. Intuitively, regions that have higher DAB also exhibit higher radial asymmetry (**Figure 5**, **third row**). This trend holds across subjects (**Figure 5**, **fourth row**), where subjects that had higher DAB in ALPS-regions also exhibited higher dispersion anisotropy.

Parallel results from the HCP-A datasets are provided in **Supplementary Material**.

### Radial asymmetry shows age-related changes, and so do geometrical features of axons

We confirm previous trends of decreasing ALPS-indices with age, where we show that globally the radial asymmetry has negative age-associations (**Figure 6**). With this, we also show that cross angle and DAB have negative age-associations, the single fiber fraction has positive age-associations. Similar trends are observed on a region-specific basis (**Figure 6**, **bottom**) where we see heterogenous changes with age in our radial asymmetry as well as heterogenous changes in geometrical features of white matter regions.

**Figure 6.**
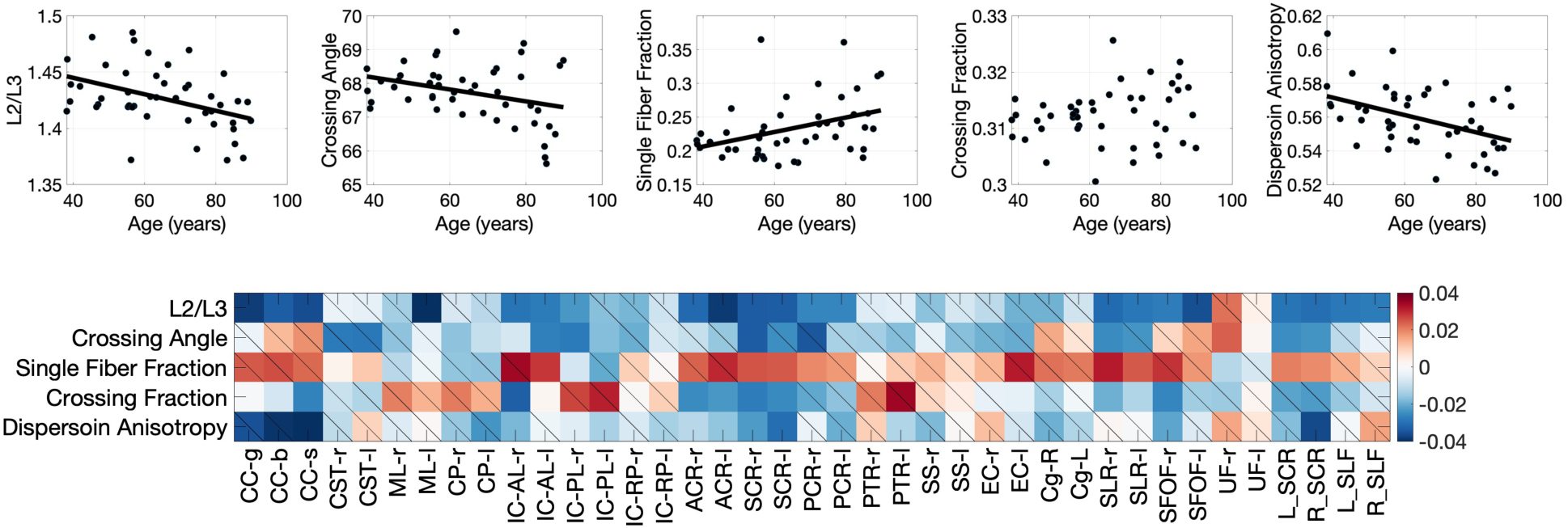
Both Radial Asymmetry and axon geometries show age-related axonal changes. (Top) all measures are plotted against age, with statistically significant associations shown as a trend-line (note no trend-line shown for non-statistically significant associations). While radial asymmetry changes with age (in agreement with ALPS-index decreases with age) so to do crossing angle, single fiber fractions, and dispersion anisotropy indices. (Bottom) these changes with age vary across pathways, shown as the beta-coefficient of the linear effects fitting (with non-significant changes shown as a diagonal line).

### Investigating PVS and WM geometrical assumptions in ALPS-related regions

**Figure 7** shows orientations for an example subject, highlighting orientation of (1) vasculature (2) primary eigenvector (V1) (3) secondary eigenvector (V2), and (4) the right-to-left orientation (RL) in the brain – shown in both axial and coronal planes. First, it is clear that our high resolution SWI and subsequent processing allow highly accurate delineation and quantification of the large medullary vein orientations, with overlaid vectors well-following the vasculature (note zoomed in region is only to highlight accurate delineation of vasculature, and does not necessarily correspond to ALPS-related regions). Second, vasculature in ALPS-regions are often orthogonal or near-orthogonal to the primary eigenvector (assumed to align with the primary direction of white matter pathways), typically within 80-90 degrees. However, the secondary eigenvector, which would be the primary driver of radial asymmetry (i.e., if PVS was influencing asymmetry), does not necessarily align with vasculature. Finally, while vasculature is indeed largely right-to-left orientation in ALPS-regions, there is large heterogeneity voxel-by-voxel, with few regions of adequate size consistently <10 degrees from the RL orientation. This heterogeneity in vasculature is observed in all subjects, and in agreement with the literature.

**Figure 7.**
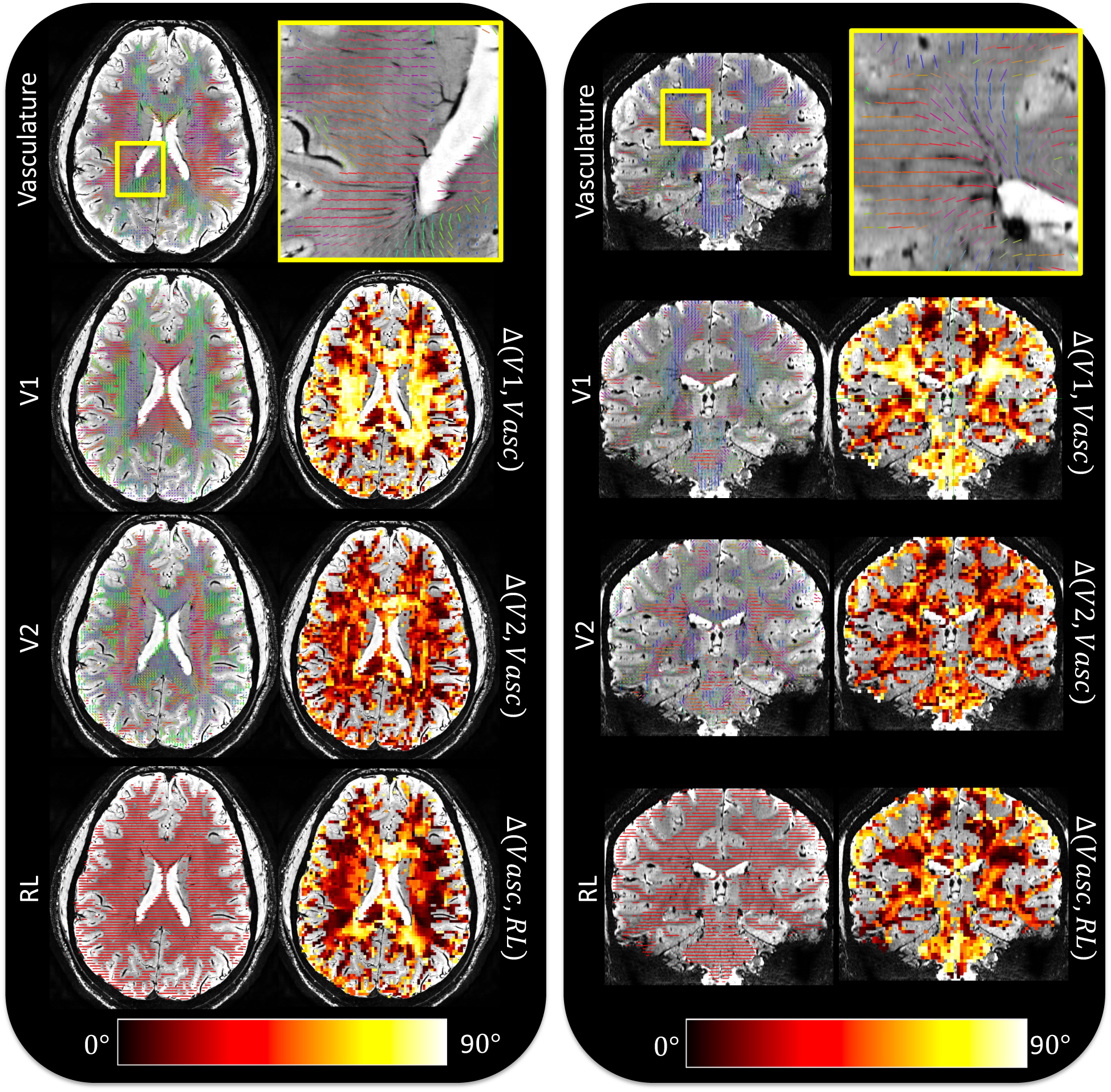
Investigating PVS and WM geometrical assumptions in ALPS-related regions. For an example subject, we investigate the relationship between vascular orientation and that from the DTI derived primary eigenvector (V1), secondary eigenvector (V2), and the right-to-left (RL) orientation. For both an axial and coronal slice, we show vascular orientation (zoom, inset), and the angular difference between vascular and white matter orientations. Note zoomed in region is only to highlight accurate delineation of vasculature, and does not necessarily correspond to ALPS-related regions. Overall, (1) vascular orientation in ALPS-associated regions are indeed near-orthogonal to WM pathways (V1), (2) but not always in alignment with the secondary eigenvector V2), (3) nor always aligned in the right-to-left orientation.

## Discussion

In this work, we investigate possible features of axonal geometry that influence the DTI-ALPS measures. Diffusivity along perivascular space (i.e. ALPS) has been proposed, and interpreted, as an indirect measure of glymphatic function. However, we show that it is potentially biased and confounded by alterations in white matter (axonal) structure, i.e., geometrical features that are prevalent throughout the entire white matter, rather than direct measures of neuro-fluid dynamics. Despite this, it is clear throughout the literature that this measure can be used to assess differences in controls and cohorts and relates to clinical and behavioral disabilities. Thus, this study aims to refine and improve the interpretation of this index and suggests the use of direct markers of fiber architecture to complement measures of diffusivity to understand biological or physiological changes occurring in pathology and disease.

### There is widespread radial asymmetry in diffusivity across white matter

We first find that radial asymmetry is widespread throughout the brain and is not confined to ALPS-related indices that exhibit this unique orthogonality of vasculature and white matter. This asymmetry exists in multiple datasets, is spatially coherent and not due to signal noise. Moreover, this asymmetry persists across a range of diffusion weightings, including high b-values typically associated with increased sensitivity to restricted diffusion within neurite-like structures, rather than the extracellular or cerebrospinal fluid-related spaces [49]. These results agree with recent work on radial asymmetry [14], and extend these results with additional datasets, additional regions, and additional diffusion weightings. Because radial asymmetry mirrors the effects that would occur to the ALPS-index due to assumed changes in PVS, together these results strongly suggest that the ALPS-index may not specifically reflect changes in PVS, but rather features of axonal geometry.

### Importantly, the ALPS-index itself is fundamentally a measure of radial diffusivity asymmetry

This relationship is illustrated in **Figure 8**, which depicts how the ALPS calculation leverages diffusivities orthogonal to dominant fiber orientations measured within association and projection regions. Mathematically, this simplifies to a ratio of diffusion tensor eigenvalues (L2/L3), where L2 reflects diffusivity aligned with the presumed perivascular orientation, and L3 reflects diffusivity orthogonal to it. The strong correlation between L2/L3 and the ALPS-index (**Figure 8**, **right**) demonstrates that index is an expression of local radial asymmetry. As such, any structural feature that alters radial diffusion asymmetry - fiber crossings, axonal undulations, dispersion - will also influence the ALPS-index. This mechanistic link serves as the central theme of this work: the ALPS-index is sensitive not only to perivascular diffusivity, but to a broader set of white matter microstructural and anatomical properties which modulate this asymmetry, and drive changes in the ALPS-index.

**Figure 8.**
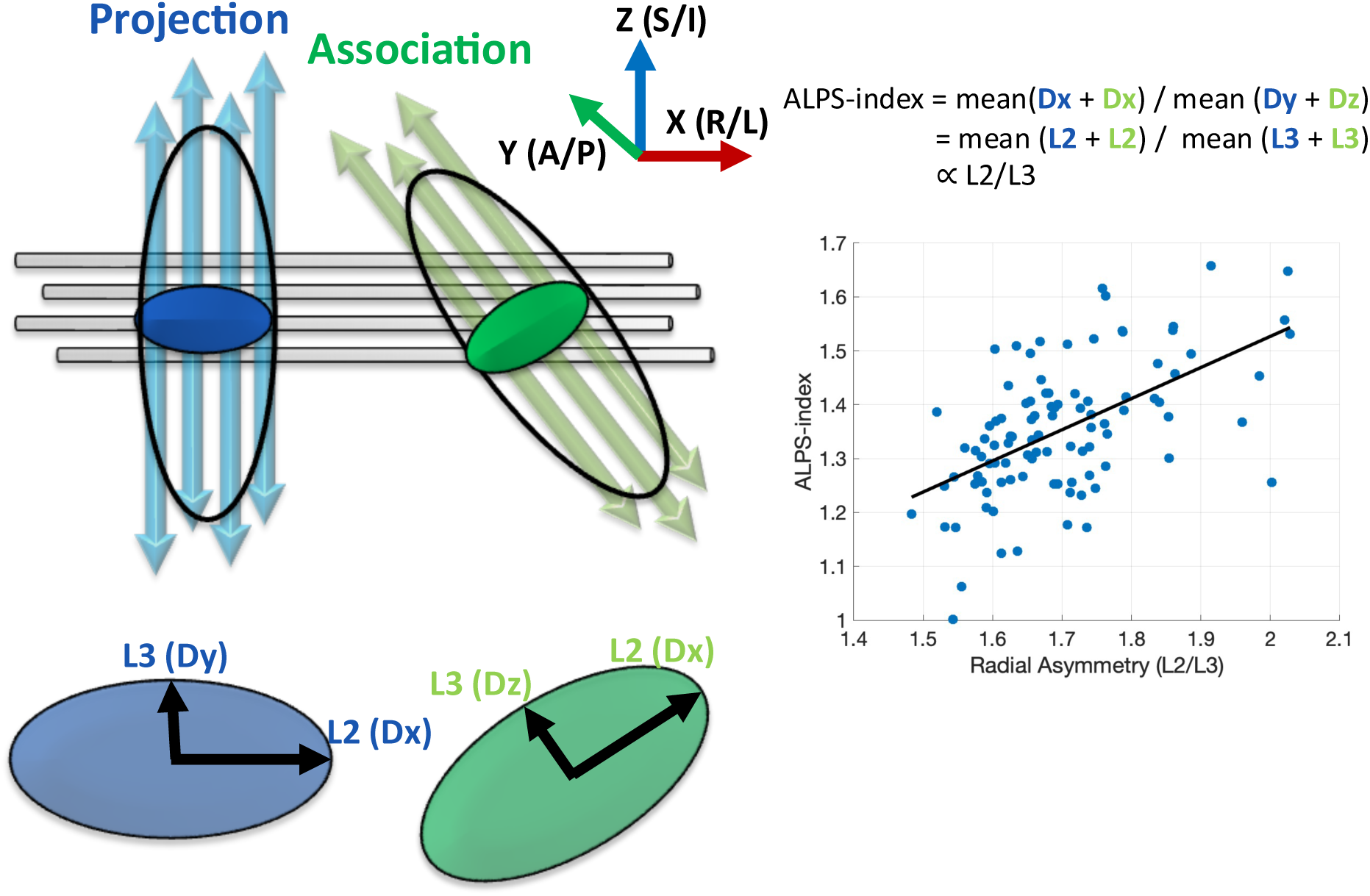
The ALPS-index is a measure of radial diffusivity asymmetry. Left: Schematic depiction of the anatomical configuration underlying the ALPS method. Medullary veins (gray cylinders) run predominantly in the right–left (x) direction, orthogonal to dominant fiber orientations of projection (blue, superior–inferior) and association (green, anterior– posterior) tracts. In these regions, diffusion tensor eigenvectors align such that λ₂ corresponds to the x-direction (Dx), and λ₃ reflects diffusivity orthogonal to both the fiber axis and perivascular direction (Dy in projection fibers, Dz in association fibers). The ALPS-index is computed as the mean of Dx across projection and association regions divided by the mean of Dy and Dz - effectively λ₂/λ₃. Right: Empirical relationship between radial asymmetry (λ₂/λ₃) and the ALPS-index across regions of interest. A strong positive association confirms that the ALPS-index is fundamentally a measure of radial asymmetry. Thus, any feature that alters radial asymmetry (investigated throughout the study), influences the ALPS index.

### Crossing fibers contribute to radial asymmetry

We next show that crossing fibers contribute to radial asymmetry and thus would contribute to a change in ALPS-index. Simulations show that axonal geometric configurations known to exist within the human brain result in asymmetries that agree with those observed in our data. Moreover, we show (in agreement with previous literature [17, 50]), that crossing fibers are widespread throughout the white matter, and specifically within regions used to derive DTI-ALPS indices. We confirm that crossing fibers (specifically the fraction of crossing fibers and angles that they cross) directly influence radial asymmetry, and thus directly influence any DTI-ALPS calculations.

Beyond this, we show that asymmetry still exists even in regions determined to contain only a single fiber population. While we know that isolating truly single fiber regions is challenging, due to multiple fiber populations traversing the same locations and the same orientations [16, 51], this suggests that there may be additional confounding factors beyond just crossing fibers.

### Dispersion and Undulation contribute to radial asymmetry

Next, we find that undulation (a waviness pattern of axonal trajectories observed in histological preparations [19, 20]) and dispersions (any geometrical effect contributing to decreased coherence) contribute to radial asymmetry. In fact, our simulations of different undulation patterns, at different diffusion weightings, and with sequence parameters matching the datasets under investigation, show radial asymmetry that well-matches what we observe in the human brain. In our real data, we find moderate to anisotropic orientation dispersion of axons, supported by improved model fitting of the anisotropic (Bingham) over isotropic (Watson) spherical distributions when fitting directly do diffusion data. The multi-compartment strategy used to model the tissue geometry specifically attributes this dispersion to the neurite compartment, thus, again this radial asymmetry is confounded by a feature of axons rather than PVS, though this interpretation is contingent on the assumptions inherent to the NODDI model[52].

This anisotropic dispersion could be due to undulations, fanning, or bending of fibers, and again is a widespread feature in the white matter of the brain. Intuitively, regions with higher modelled DAB have higher radial asymmetry, and individuals with higher DAB exhibit higher radial asymmetry in ALPS-regions. While again the radial asymmetry is confounded by a geometrical feature of white matter, this anisotropic orientation dispersion can serve as a useful marker of pathology, reflecting subtle changes in dispersion and coherence.

### Age-related axonal changes

Several studies using the DTI-ALPS method have investigated the association between age and the ALPS-index in both health and disease [7, 8, 12, 13]. In general, they find negative associations between ALPS-indices and aging, particularly for cohorts >40. In agreement with this, we find age-related changes in radial asymmetry. In parallel with this, we show that these confounding factors (crossing fibers, axonal dispersion) and their associated indices show similar trends. Interestingly, these trends are not the same across all pathways, with varying degrees of geometrical change depending on location and pathway type (commissural, association, projection). In this study, we don’t intend to interpret the biophysical aging mechanisms associated with potentially reduced crossing fibers (axonal degeneration) or decreased dispersion anisotropy, but use this demonstration to highlight that observed ALPS-index changes are again associated with geometrical changes in aging.

### Orientation of PVS and WM

DTI-ALPS takes advantage of the unique tissue geometry adjacent to the lateral ventricles, where it is assumed that the vasculature runs left-to-right while pathways run both superior-inferior (projection pathways of the SCR) and anterior-posterior (association pathways of the SLF). By measuring the ratio of radial asymmetry in each pathway, in theory, DTI-ALPS isolates the left-right diffusivity attributed to PVS. We find that this arrangement, again in agreement with the literature, is indeed generally true – left-right orientation of vasculature, superior-inferior orientation of projection pathways, and anterior-to-posterior orientation of association pathways. However, due to heterogeneity in both tissue types, there is no large region in which all of these anatomical and diffusional assumptions - orthogonal vein and fiber orientation, single-fiber geometry, and consistent eigenvector alignment - hold true (within a reasonable angular error) and are free of crossing fibers. For example, while predominantly left-right throughout the entire SCR + SLF area, vasculature can vary by as much as ∼30 degrees from this orientation, depending on exact location, again decreasing the specificity of ALPS-indices to diffusivity attributed to vasculature. Further, if the main contributor to axial asymmetry was diffusion along perivascular space, then it is intuitive that the secondary eigenvector would also point along the vascular orientation, which in general is not true. These results highlight challenges with choosing regions of interest for ALPS-calculation, and possible effects that contribute to decreased specificity (i.e., region placement resulting in non-left-right oriented vasculature or non-orthogonal white matter).

### Perivascular space size and shape and diffusion sensitivity

These results lead us to ask what diffusion might be able to tell us about PVS. Recent quantifications in murine pial arteries give us an idea of the size and shape of PVS [15]. First, these spaces exhibit considerable variability, with cross sectional geometries not easily described by cylindrical nor ellipsoidal geometries, instead better characterized as a complex polynomial or spline functions surrounding a central cylindrical vessel, with the width/height of the PVS decaying as a function of distance from the vessel (see Figure 10 from [15]). The PVS measured in [15] was associated with vessels with radii ranging from 19-29um (25^th^-75^th^ percentiles), and had spaces with heights ranging from 24-45um (25^th^-75^th^ percentiles) immediately adjacent to the vessels. Overall, these are much larger structures than the vessels themselves, and an order of magnitude larger than axons and axonal spacing – which result in low resistance and facilitation of fluid movement. This means, however, that moderate-to-high b-values (both typical DTI studies with b∼1000s/mm2, and the current study reaching b=3000s/mm2) will have significant signal attenuation in these spaces and not be highly specific to PVS. However, it should be note that these measurements in the mouse model does not preclude smaller PVS that may not have been measured with their optical imaging.

The complexity and microscopic dimensions of PVS thus presents challenges for their specific detection and quantification using diffusion MRI. Indeed, Sepehrband et al. [53] demonstrated that even a small fractional contribution from anisotropic perivascular fluid can markedly bias DTI metrics, artificially elevating MD and reducing FA, thereby confounding interpretations that traditionally attribute such changes solely to tissue. Optimistically, this means that our typical sequences are indeed sensitive in some way to PVS. As proposed in [53], incorporating multi-shell diffusion sequences (with low b-values) coupled with multi-compartment modeling may be capable of explicitly distinguishing anisotropic fluid compartments from the surrounding tissue – however, this depends on the diffusion and size/shape characteristics of the PVS. This would allow characterization of distinct diffusion profiles for both parenchymal tissue and perivascular fluid (with higher, nearly free water, diffusivity). Such approaches are essential for disentangling true microstructural tissue alterations from geometrically induced fluid compartment biases, thus improving specificity in interpreting diffusion MRI biomarkers in both clinical and research settings.

To further contextualize the sensitivity of diffusion MRI to perivascular diffusion, and explore whether a genuine PVS signal could influence the asymmetry metrics used in ALPS, we performed a simplified supplementary simulation incorporating an asymmetric PVS compartment with a volume fraction of 10% and diffusivities consistent with prior experimental estimates [53]. Simulated signal decay curves demonstrate that at high b-values (e.g., b = 3000 s/mm²), the asymmetric contribution from the PVS compartment is effectively nulled due to its rapid apparent diffusivity. At b = 1000 s/mm² - more typical of clinical DTI protocols - PVS signal remains detectable but induces only minimal changes in derived radial asymmetry measures. These findings reinforce that the ALPS-index is largely insensitive to true perivascular diffusion effects under standard acquisition parameters. Even if no structural confounds were present (e.g., crossing fibers or axonal undulations), the ALPS-index itself would still exhibit limited sensitivity to PVS-specific diffusion. Supporting simulation details and quantitative model outputs are shown in **Supplementary Experiment 1**. We do not include this within primary results, as our study is largely focused on potential confounds that may drive asymmetry (rather than the sensitivity of the diffusion signal itself), and we do not claim that this simulation captures the true diffusivity of each compartment (for which more validation is needed).

### Limitations

The current study has several limitations. First, we used only two cross-sectional datasets, that are not typical of clinically acquired data available on most 1.5T or 3T scanners. However, these results represent a “best case” scenario in terms of confounds and biases because of their high spatial resolution and limited partial volume effects (i.e. crossing fiber effects will become more prevalent with larger voxels). Second, we investigated only well-known geometrical effects studied in the diffusion community – crossing fibers and dispersion/undulations. Additional geometrical configurations that may introduce radial asymmetries including larger scale fanning and branching (observable with asymmetry fiber orientation distributions), and it is also possible that sheet-like geometry, where fiber pathways cross on 2D (curved) surfaces, observable in some brain regions may create anisotropic structure on larger scales contributing to asymmetry[54, 55]. Both are potentially interesting geometrical features that have been under-investigated.

## Conclusion

This study demonstrates that diffusion tensor imaging along perivascular spaces (DTI-ALPS) measures are influenced by widespread radial asymmetry arising from crossing fibers, and axonal undulations and dispersion in white matter. Our findings indicate that radial asymmetry, and hence DTI-ALPS metrics, are not exclusively reflective of perivascular diffusion, but rather significantly confounded by underlying axonal geometry. Interpretations of ALPS-derived metrics as biomarkers of glymphatic function must carefully consider these anatomical and microstructural complexities, and future studies should model this compartment with tailored sequences and modelling strategies.

## Acknowledgements

This work was supported by the National Institutes of Health (NIH) under award numbers K01EB032898 (KS), R01EB017230 (BL). CMWT is supported by the Wellcome Trust [215944/Z/19/Z]. For the purpose of open access, the author has applied a CC BY public copyright license to any Author Accepted Manuscript version arising from this submission.

## Author Contributions

KGS, CT, AN, MC, AA, BL, MD

## Ethics

All participants from whom data were used in this manuscript, provided written informed consent (and consent to publish) according to the declaration of Helsinki.

## Data and Code Availability

## Supplementary Experiments

**Supplementary Experiment 1.**
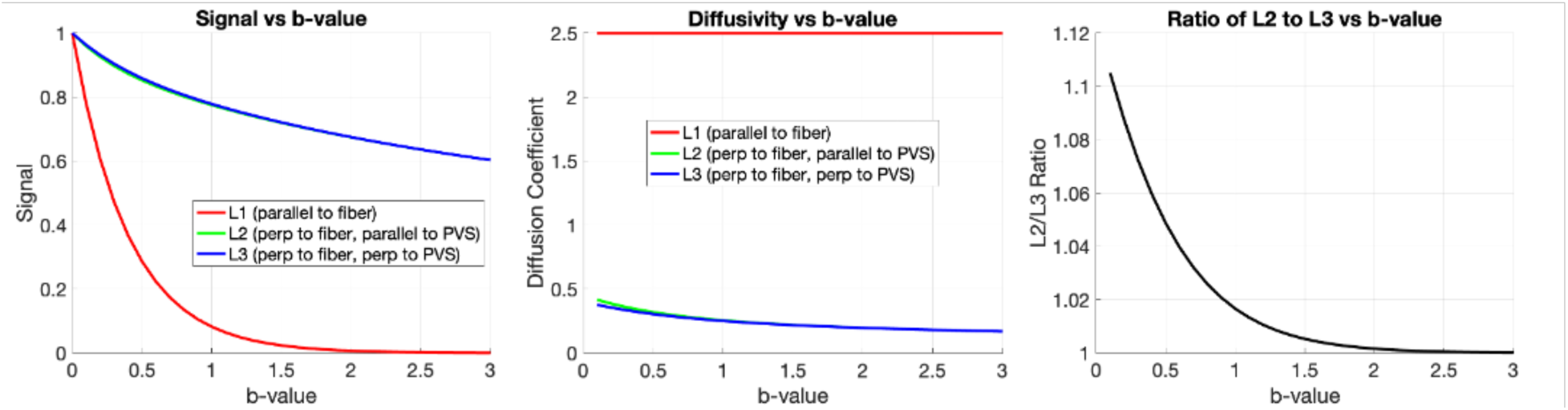
Investigating how a true PVS/APLS effect influences our metrics of asymmetry, i.e., would a true ALPS-related changes influence the asymmetry index? A simple 3 compartment model was simulated, incorporating an asymmetric PVS compartment with a volume fraction of 10% and diffusivities consistent with prior experimental estimates [53] (along perivascular space diffusivity=3, radial to perivascular space diffusivity=2.5), with an intra-axonal compartment (axial diffusivity = 2.5, radial diffusivity = 0), and an extracellular compartment (isotropic diffusivity = 2.5). The signal was simulated across a range of b-values for (1) along the WM fiber and perpendicular to PVS (L1) (2) perpendicular to the fiber and parallel to PVS (L2), and (3) perpendicular to the fiber and perpendicular to PVS (L3): S1 = 0.1*exp(-b*2.5) + 0.4*exp(-b*2.5) + 0.5*exp(-b*2.5); S2 = 0.1*exp(-b*3) + 0.4*exp(-b*0) + 0.5*exp(-b*0.3); S3 = 0.1*exp(-b*2.5) + 0.4*exp(-b*0) + 0.5*exp(-b*0.3); From this, the diffusivities are calculated along each direction, and the ratio of L2 to L3 (i.e., the asymmetry index) was derived. With these assumed diffusivities, high b-values are not sensitive to asymmetric PVS effects.

## References

1. Wardlaw, J.M., et al., Perivascular spaces in the brain: anatomy, physiology and pathology. Nat Rev Neurol, 2020. 16(3): p. 137–153.

2. Iliff, J.J., et al., A paravascular pathway facilitates CSF flow through the brain parenchyma and the clearance of interstitial solutes, including amyloid beta. Sci Transl Med, 2012. 4(147): p. 147ra111.

3. Buccellato, F.R., et al., The Role of Glymphatic System in Alzheimer’s and Parkinson’s Disease Pathogenesis. Biomedicines, 2022. 10(9).

4. Taoka, T., et al., Evaluation of glymphatic system activity with the diffusion MR technique: diffusion tensor image analysis along the perivascular space (DTI-ALPS) in Alzheimer’s disease cases. Jpn J Radiol, 2017. 35(4): p. 172–178.

5. Taoka, T., et al., Diffusion Tensor Image Analysis ALong the Perivascular Space (DTI-ALPS): Revisiting the Meaning and Significance of the Method. Magn Reson Med Sci, 2024. 23(3): p. 268–290.

6. Chang, H.I., et al., Gray matter reserve determines glymphatic system function in young-onset Alzheimer’s disease: Evidenced by DTI-ALPS and compared with age-matched controls. Psychiatry Clin Neurosci, 2023. 77(7): p. 401–409.

7. Steward, C.E., et al., Assessment of the DTI-ALPS Parameter Along the Perivascular Space in Older Adults at Risk of Dementia. J Neuroimaging, 2021. 31(3): p. 569–578.

8. McKnight, C.D., et al., Diffusion along perivascular spaces reveals evidence supportive of glymphatic function impairment in Parkinson disease. Parkinsonism Relat Disord, 2021. 89: p. 98–104.

9. Ma, X., et al., Diffusion Tensor Imaging Along the Perivascular Space Index in Different Stages of Parkinson’s Disease. Front Aging Neurosci, 2021. 13: p. 773951.

10. Carotenuto, A., et al., Glymphatic system impairment in multiple sclerosis: relation with brain damage and disability. Brain, 2022. 145(8): p. 2785–2795.

11. Hagiwara, A., et al., Glymphatic System Dysfunction in Myelin Oligodendrocyte Glycoprotein Immunoglobulin G Antibody-Associated Disorders: Association with Clinical Disability. AJNR Am J Neuroradiol, 2023. 45(1): p. 66–71.

12. Matsushita, S., et al., The Association of Metabolic Brain MRI, Amyloid PET, and Clinical Factors: A Study of Alzheimer’s Disease and Normal Controls From the Open Access Series of Imaging Studies Dataset. J Magn Reson Imaging, 2024. 59(4): p. 1341–1348.

13. Zhang, W., et al., Glymphatic clearance function in patients with cerebral small vessel disease. Neuroimage, 2021. 238: p. 118257.

14. Wright, A.M., et al., Exploring Radial Asymmetry in MR Diffusion Tensor Imaging and Its Impact on the Interpretation of Glymphatic Mechanisms. J Magn Reson Imaging, 2024. 60(4): p. 1432–1441.

15. Raicevic, N., et al., Sizes and shapes of perivascular spaces surrounding murine pial arteries. Fluids Barriers CNS, 2023. 20(1): p. 56.

16. Schilling, K.G., et al., Prevalence of white matter pathways coming into a single white matter voxel orientation: The bottleneck issue in tractography. Hum Brain Mapp, 2021.

17. Jeurissen, B., et al., Investigating the prevalence of complex fiber configurations in white matter tissue with diffusion magnetic resonance imaging. Hum Brain Mapp, 2013. 34(11): p. 2747–66.

18. Georgiopoulos, C., et al., Diffusion tensor imaging along the perivascular space: the bias from crossing fibres. Brain Commun, 2024. 6(6): p. fcae421.

19. Budde, M.D. and J. Annese, Quantification of anisotropy and fiber orientation in human brain histological sections. Front Integr Neurosci, 2013. 7: p. 3.

20. Schilling, K.G., et al., Histological validation of diffusion MRI fiber orientation distributions and dispersion. Neuroimage, 2018. 165: p. 200–221.

21. Jeffery, G., PNS features of rodent optic nerve axons. J Comp Neurol, 1996. 366(2): p. 370–8.

22. Lontis, E.R., K. Nielsen, and J.J. Struijk, In vitro magnetic stimulation of pig phrenic nerve with transverse and longitudinal induced electric fields: analysis of the stimulation site. IEEE Trans Biomed Eng, 2009. 56(2): p. 500–12.

23. Sunderland, S. and K.C. Bradley, STRESS-STRAIN PHENOMENA IN HUMAN PERIPHERAL NERVE TRUNKS1. Brain, 1961. 84(1): p. 102–119.

24. Lee, H.H., et al., A time-dependent diffusion MRI signature of axon caliber variations and beading. Commun Biol, 2020. 3(1): p. 354.

25. Brabec, J., S. Lasic, and M. Nilsson, Time-dependent diffusion in undulating thin fibers: Impact on axon diameter estimation. NMR Biomed, 2020. 33(3): p. e4187.

26. Nilsson, M., et al., The role of tissue microstructure and water exchange in biophysical modelling of diffusion in white matter. MAGMA, 2013. 26(4): p. 345–70.

27. Nilsson, M., et al., The importance of axonal undulation in diffusion MR measurements: a Monte Carlo simulation study. NMR Biomed, 2012. 25(5): p. 795–805.

28. Lee, H.H., et al., The impact of realistic axonal shape on axon diameter estimation using diffusion MRI. Neuroimage, 2020. 223: p. 117228.

29. Bernier, M., S.C. Cunnane, and K. Whittingstall, The morphology of the human cerebrovascular system. Hum Brain Mapp, 2018. 39(12): p. 4962–4975.

30. Nonaka, H., et al., Microvasculature of the human cerebral white matter: arteries of the deep white matter. Neuropathology, 2003. 23(2): p. 111–8.

31. Bernier, M., et al. Human cerebral white-matter vasculature imaged using the blood-Pool contrast agent ferumoxytol: bundle-specific vessels and vascular density. in Proc Intl Soc Mag Reson Med. 2020.

32. Smirnov, M., C. Destrieux, and I.L. Maldonado, Cerebral white matter vasculature: still uncharted? Brain, 2021. 144(12): p. 3561–3575.

33. Ruiz, D.S., H. Yilmaz, and P. Gailloud, Cerebral developmental venous anomalies: current concepts. Ann Neurol, 2009. 66(3): p. 271–83.

34. Van Essen, D.C., et al., The Human Connectome Project: a data acquisition perspective. Neuroimage, 2012. 62(4): p. 2222–31.

35. Glasser, M.F., et al., The minimal preprocessing pipelines for the Human Connectome Project. Neuroimage, 2013. 80: p. 105–24.

36. Basser, P.J., J. Mattiello, and D. LeBihan, MR diffusion tensor spectroscopy and imaging. Biophys J, 1994. 66(1): p. 259–67.

37. Pierpaoli, C., et al., Diffusion tensor MR imaging of the human brain. Radiology, 1996. 201(3): p. 637–48.

38. Tournier, J.D., et al., MRtrix3: A fast, flexible and open software framework for medical image processing and visualisation. Neuroimage, 2019. 202: p. 116137.

39. Mori, S., K. Oishi, and A.V. Faria, White matter atlases based on diffusion tensor imaging. Curr Opin Neurol, 2009. 22(4): p. 362–9.

40. Liu, X., et al., Cross-Vendor Test-Retest Validation of Diffusion Tensor Image Analysis along the Perivascular Space (DTI-ALPS) for Evaluating Glymphatic System Function. Aging Dis, 2024. 15(4): p. 1885–1898.

41. Daducci, A., et al., Quantitative comparison of reconstruction methods for intra-voxel fiber recovery from diffusion MRI. IEEE Trans Med Imaging, 2014. 33(2): p. 384–99.

42. Dhollander, T., D. Raffelt, and A. Connelly. Unsupervised 3-tissue response function estimation from single-shell or multi-shell diffusion MR data without a co-registered T1 image. in ISMRM workshop on breaking the barriers of diffusion MRI. 2016. Lisbon, Portugal.

43. Dhollander, T. and A. Connelly, A novel iterative approach to reap the benefits of multi-tissue CSD from just single-shell (+b=0) diffusion MRI data. 2016.

44. Tariq, M., et al., Bingham-NODDI: Mapping anisotropic orientation dispersion of neurites using diffusion MRI. Neuroimage, 2016. 133: p. 207–223.

45. Veraart, J., E. Fieremans, and D.S. Novikov, Diffusion MRI noise mapping using random matrix theory. Magn Reson Med, 2016. 76(5): p. 1582–1593.

46. Fadnavis, S., J. Batson, and E. Garyfallidis, Patch2Self: denoising diffusion MRI with self-supervised learning. arXiv preprint arXiv:2011.01355, 2020.

47. Andersson, J.L., S. Skare, and J. Ashburner, How to correct susceptibility distortions in spin-echo echo-planar images: application to diffusion tensor imaging. Neuroimage, 2003. 20(2): p. 870–88.

48. Cai, L.Y., et al., PreQual: An automated pipeline for integrated preprocessing and quality assurance of diffusion weighted MRI images. Magn Reson Med, 2021. 86(1): p. 456–470.

49. Novikov, D.S., V.G. Kiselev, and S.N. Jespersen, On modeling. Magn Reson Med, 2018. 79(6): p. 3172–3193.

50. Schilling, K., et al., Can increased spatial resolution solve the crossing fiber problem for diffusion MRI? NMR Biomed, 2017. 30(12).

51. Maier-Hein, K.H., et al., The challenge of mapping the human connectome based on diffusion tractography. Nat Commun, 2017. 8(1): p. 1349.

52. Novikov, D.S., et al., Quantifying brain microstructure with diffusion MRI: Theory and parameter estimation. NMR Biomed, 2018: p. e3998.

53. Sepehrband, F., et al., Perivascular space fluid contributes to diffusion tensor imaging changes in white matter. Neuroimage, 2019. 197: p. 243–254.

54. Tax, C.M.W., et al., Sheet Probability Index (SPI): Characterizing the geometrical organization of the white matter with diffusion MRI. Neuroimage, 2016. 142: p. 260–279.

55. Tax, C.M.W., et al., Quantifying the brain’s sheet structure with normalized convolution. Med Image Anal, 2017. 39: p. 162–177.

