## Supplementary Material for "White matter geometry confounds Diffusion Tensor Imaging Along Perivascular Space (DTI-ALPS) measures"

Keywords: ALPS; crossing fibers; DTI-ALPS; Undulation; Dispersion; Glymphatic System

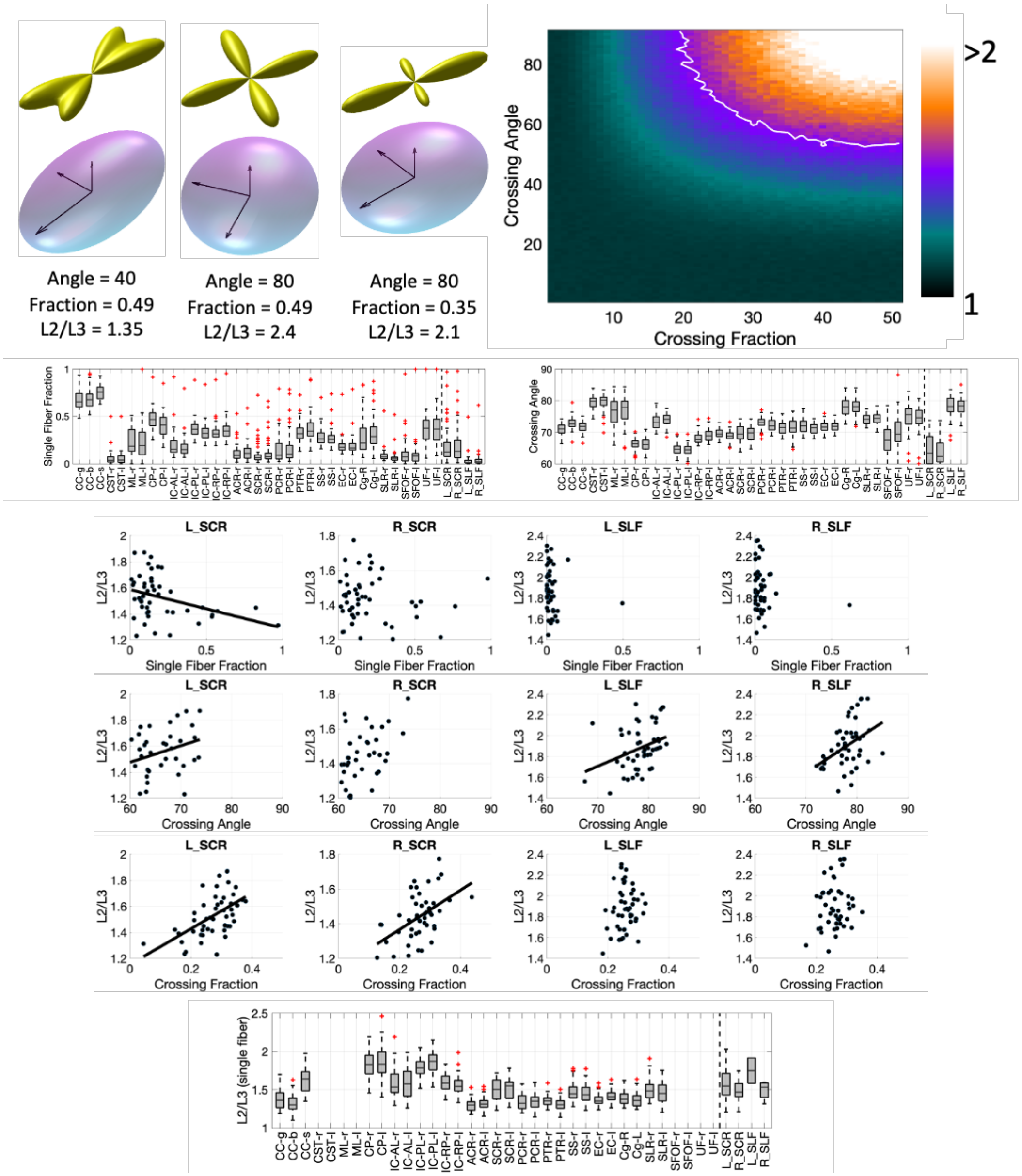


**Supplementary Figure 1**. (In parallel to main manuscript Figure 3 – Supplementary Figure 1 based on HCP-Aging data, main figure 3 based on HCP Young Adult Data). Most white matter regions contain crossing fibers, which contribute to radial asymmetry. (First Row) crossing fibers at different crossing angles and fiber fractions, resulting in radial asymmetry are visualized, as well as simulation results showing that increasing crossing angle and crossing fraction result in increased radial asymmetry. (Second Row) Single Fiber Fraction and Crossing Angle are shown averaged in each white matter region. (Middle) Radial asymmetry is plotted against Single Fiber Fraction, Crossing Angle, and Crossing Fraction for ALPS-specific ROIs, across subjects, confirming that crossing fibers significantly influence asymmetry measures. (Bottom) Axial asymmetry remains even in single fiber voxels within each region.

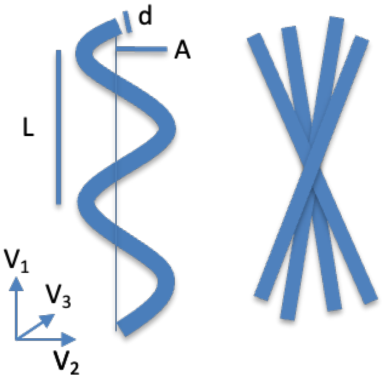


| d | L | A | delta | DELTA | bval | asym | noise |
|---|---|---|---|---|---|---|---|
| 1 | 24 | 4 | 10.6 | 43.1 | 1000 | 1.35 | 1.36 |
| 1 | 300 | 50 | 10.6 | 43.1 | 1000 | 2.07 | 2.06 |
| 1 | 24 | 4 | 10.6 | 43.1 | 2000 | 1.9 | 1.39 |
| 1 | 300 | 50 | 10.6 | 43.1 | 2000 | 2.44 | 2.32 |
| 1 | 24 | 4 | 10.6 | 43.1 | 3000 | 1.52 | 1.48 |
| 1 | 300 | 50 | 10.6 | 43.1 | 3000 | 2.90 | 2.43 |

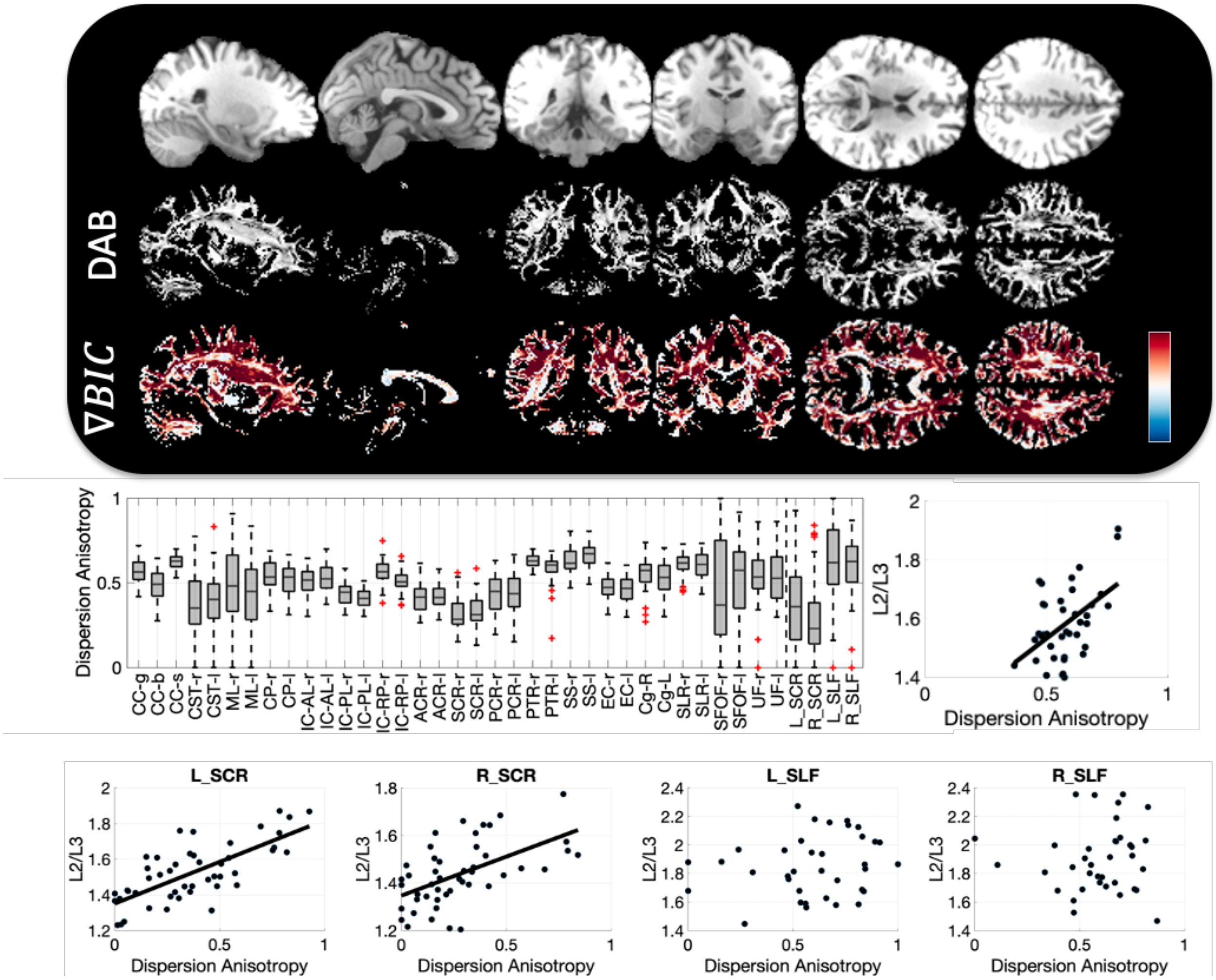


**Supplementary Figure 2.** (Parallel to Figure 5 of the main text – where Supplementary Figure 2 shows results for HCPA, main figure 5 shows results for HCP Young Adult data). Dispersion and Undulation contribute to radial asymmetry. (Top) example undulation and dispersion cartoons, highlighting within-plane orientation dispersion. Simulation results with two substrates show that undulations can contribute to radial asymmetry on par with that observed in empirical data. (Second Row) Example sagittal, coronal, and axial slices showing the Dispersion Anisotropy Index (DAB) (gray-scale) and the difference in BIC between Watson and Bingham distributions (Red indicates preference for Bingham, Blue indicates preference for Watson distribution). (Third Row) Dispersion Anisotropy (DAB) across all white matter regions; radial asymmetry plotted against Dispersion Anisotropy (DAB) across all regions shows strong relationship between these two measures of asymmetry. (Bottom Row). Similarly, Radial Asymmetry plotted against DAB across subjects shows strong relationships between these two measures in ALPS-specific ROIs, with a trend-line shown for regions with statistically significant associations.

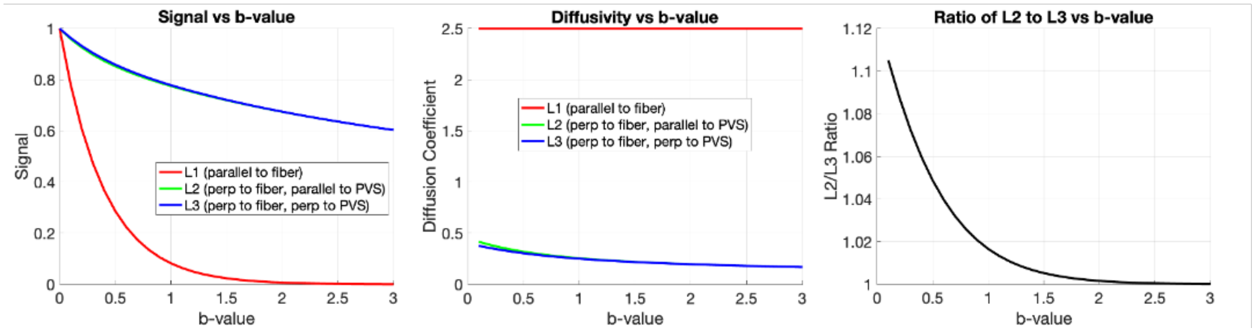


**Supplementary Experiment 1.** Investigating how a true PVS/APLS effect influences our metrics of asymmetry, i.e., would a true ALPS-related changes influence the asymmetry index? A simple 3 compartment model was simulated, incorporating an asymmetric PVS compartment with a volume fraction of 10% and diffusivities consistent with prior experimental estimates [53] (along perivascular space diffusivity=3, radial to perivascular space diffusivity=2.5), with an intra-axonal compartment (axial diffusivity = 2.5, radial diffusivity = 0), and an extracellular compartment (isotropic diffusivity = 2.5). The signal was simulated across a range of b-values for (1) along the WM fiber and perpendicular to PVS (L1) (2) perpendicular to the fiber and parallel to PVS (L2), and (3) perpendicular to the fiber and perpendicular to PVS (L3):

```
S1 = 0.1*exp(-b*2.5) + 0.4*exp(-b*2.5) + 0.5*exp(-b*2.5);
S2 = 0.1*exp(-b*3)   + 0.4*exp(-b*0)   + 0.5*exp(-b*0.3);
S3 = 0.1*exp(-b*2.5) + 0.4*exp(-b*0)   + 0.5*exp(-b*0.3);
```

From this, the diffusivities are calculated along each direction, and the ratio of L2 to L3 (i.e., the asymmetry index) was derived. With these assumed diffusivities, high b-values are not sensitive to asymmetric PVS effects.
